# Inositolphosphate glycans accumulate and suppress plant defense during Arabidopsis/Botrytis interaction

**DOI:** 10.1101/2023.11.16.567443

**Authors:** Luka Lelas, Justine Rouffet, Alexis Filachet, Julien Sechet, Elodie Akary, Thierry Desprez, Samantha Vernhettes, Aline Voxeur

## Abstract

This study investigates the presence and significance of previously undiscovered oligosaccharides that accumulate during the interaction between *Arabidopsis thaliana* and *Botrytis cinerea*, a pathogenic fungus. Initially focused on characterizing cell wall-derived oligosaccharides, the research uncovered inositol phosphate glycans (IPGs) originating from plant sphingolipids, specifically glycosylinositol phosphorylceramides. Advanced chromatography, mass spectrometry techniques and molecular biology were employed to identify these IPGs, determine their origins, and study their role in the *A. thaliana-B. cinerea* interaction. Contrary to the conventional belief that oligosaccharides trigger plant defense, this research suggests that *B. cinerea* releases IPGs identical to those generated by host plant to actually downregulate plant defense mechanisms. This discovery offers insight into the dynamic strategies used by *B. cinerea* to evade plant defenses and establish successful infections.

**One-Sentence Summary:** A plant pathogen releases products identical to those generated by host plant that aids in evasion of the plant defense.

## Main Text

*Botrytis cinerea*, commonly known as gray mold fungus, is a plant necrotrophic pathogen that can infect a wide range of plant species. This fungus is particularly notorious for its feeding on the plant cell walls, an interaction that raises critical questions about its biological mechanisms and the plant’s defense responses (*1*). Elucidating the intricacies of *B. cinerea’*s biology and its interactions with host plant helps in the development of sustainable and effective strategies to mitigate its impact on agriculture, benefiting both the scientific community and society at large.

Oligosaccharins are biological active oligosaccharides that can be derived from plant cell walls and be produced upon plant pathogen interaction (*2*). They are known for their diverse effects in plants, including promoting growth in low- and normal-light conditions (*3-7*) and anti-auxin activity (*8,9*). Moreover, while we know for decades that oligogalacturonides (OGs), α-1,4-linked galacturonic acid oligomers deriving from pectins, trigger plant defense responses upon exogenous application (*10-12*), the detection and the structural characterization of these oligosaccharins within plants infected by *B. cinerea* has been a recent development (*13*).

In our study, we have uncovered previously unknown oligosaccharides that accumulate during the interaction between *Arabidopsis thaliana* and *B. cinerea*. In contrast to our initial expectations of characterizing cell wall-derived oligosaccharides, we have identified a substantial presence of inositol phosphate glycans (IPGs) which become highly abundant during infection and originate from vital and intricate lipids of plant cell membranes: the glycosylinositol phosphorylceramides (GIPCs),

### Unknown oligosaccharides accumulate during *A. thaliana/B. cinerea* interaction

To begin, we employed high-performance size-exclusion chromatography (HP-SEC) in conjunction with a high-resolution mass spectrometry (HRMS) method in negative mode to establish a systematic approach for identifying potential oligosaccharides that accumulate during infection over time. We focused on ions consistently increasing between 12- and 18-hours post-infection, possessing a mass-to-charge ratio (m/z) exceeding 263 and a retention time below 8 minutes, indicative of a dipentose (**Fig. 1A, Dataset S1**). This analysis led to the identification of 36 candidates, which included 16 candidates with predicted formulae and retention time compatible with OGs we have previously identified (*13*), four of which were likely oxidized probably through plant OG oxidases (*14*) while 12 were unsaturated (-H_2_O) and likely result from pectin lyase activity (*13*) (**Fig. 1B**). Out of the remaining 20 candidates, seven were excluded due to the unreliable elemental compositions. Consequently, our methodology led to the recognition of 13 candidates, with nine being phosphorylated, while three lacked elemental compositions compatible with an oligosaccharide structure (**Fig. 1C**). Remarkably, eight of these candidates were also detected in the cell wall water extract of non-infected plants.

**Figure 1.**
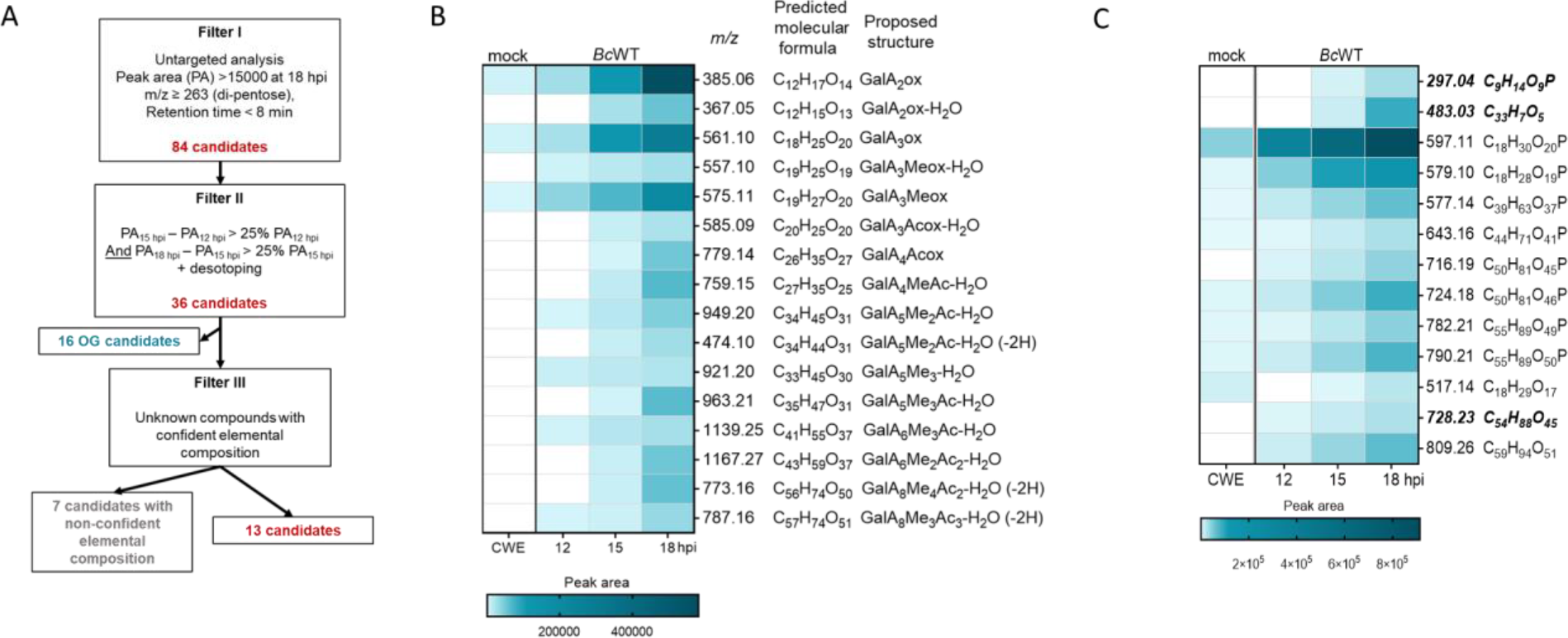
Candidate selection using a three-step filtering strategy. **(A)** Flowchart illustrating of the three-step filtering strategy employed to select potential oligosaccharides that accumulate during infection. **(B**) Heat map displaying the putative oligogalacturonides (OG) accumulated over time during the infection. **(C**) Heat map showing the non-OG candidates that accumulate over time during the infection. Candidates in bold and italic lack elemental compositions compatible with an oligosaccharide structure. PA: Peak area; hpi: hours post infection; OG: oligogalacturonides; CWE: Cell wall water extract; BcWT: *B. cinerea* wild type strain; GalA: Galacturonic acid; Me: Methylester group; Ac: Acetylester group; OGs are named GalA_x_Me_y_Ac_z_. Subscript numbers indicate the DP and the number of methyl- and acetylester groups, respectively.

### Acetylated xyloglucan-derived oligosaccharides and inositol phosphate glycans are produced during *A. thaliana/B. cinerea* interaction

The candidate with the largest molecular formula at *m/z* 809 is doubly charged (predicted formula C_60_H_98_O_50_) was detected as a formic acid adduct [*M*+HCOO^-^-H]^2-^. Its fragmentation resulted in the formation of its [*M*-H]^-^ form at *m/z* 1573, along with ions at *m/z* 161 (anhydro-hexosyl unit), 1411 [M-C_6_H_10_O_5_-H]^-^and 1279 [M-C_6_H_10_O_5_-C_5_H_8_O_4_-H]^-^ suggesting the presence of hexosyl and pentosyl residues (**Fig. S1**). We attributed the doubly charged ion at *m/z* 726, produced via a 120-Da loss, to a ^2,4^A_n_ cross-ring cleavage in a C_4_-substituted hexose. Ions at 675 [M-Hex-C_2_H_4_O_2_-2H]^2-^and 666 [M-Hex-C_2_H_4_O_2_-H_2_O-2H]^2-^ were assigned to a ^0,2^A_n-1_ cross-ring cleavage which is diagnostic of C_4_- and C_6_-substituted hexose. The prominent and smaller single charged fragments at *m/z* 643 and 455 were accompanied by ions at *m/z* 541 and 353, respectively, corresponding to a 102-Da neutral loss. This MS^2^ pattern is characteristic of xyloglucan oligosaccharide fragmentations known for their prominent D-type fragment ions. These fragment result from double cleavage and correspond to entire inner xyloglucan side chains (*15*). They produce ^0,4^A_i_ cross-ring cleavage ions, diagnostic for the presence of (1,6)-linkage, via a 102-Da loss from anhydro-glycosyl unit. Consequently, we attributed the D-type ion at *m/z* 455 and its ^0,4^A_β_ corresponding ion at *m/z* 353 to the presence of Galactose-Xylose-Glucose block (represented by the letter L). Additionally, *m/z* 643 (D), 541 (^0,4^A_iα_) and 499 (^0,4^A_iα_ -Acetyl) were attributed to the presence of an acetylated Fucose-Galactose-Xylose-Glucose block (represented by the letter F). Based on these results and the well-known structure of xyloglucan, we concluded that the candidate at *m/z* 809 is an acetylated XLFG oligosaccharide.

We next focused our study on the most abundant candidate at m/z 597 [*M* - H]^-^ for which the software predicted the formula C_18_H_30_O_20_P^-^. This compound coelutes with the m/z 579 ion (predicted formula C_18_H_29_O_19_P^-^) which we attributed to the formation of a [*M*-H-H2O]^-^ anion. The *m/z* 597 MS^2^ spectrum displays a *m/z* 97 ion corresponding to H_2_PO_4-_that comforts the predicted formula (**Fig. 2A**). Furthermore, the *m/z* 259 and 241 ions are diagnostic of phosphorylated inositol (Ino-P), they are notably found in MALDI-MS/MS analysis of GIPCs (*16, 17*). The ion at *m/z* 241 was next assigned the loss of an uronic acid (UA) from the *m/z* 373 ion that has been shown to represent a useful signature of the GIPC polar head (*16, 17)*. It is likely formed from the decarboxylation of the *m/z* 417 ion, the latter corresponding to the loss of a hexose (Hex) moiety from the parental ion [*M*-H-180]^-^. Altogether, this fragmentation pattern suggests that *m/z* 597 ion corresponds to an inositolphosphate glycan (IPGs) composed of a phosphorylated inositol, an uronic acid and a hexose that might originate from *A. thaliana* Series A GIPCs. We next fragmented the candidate at *m/z* 517 (C_18_H_29_O_17-_) (**Fig. 2B**). We observed fragment ions at *m/z* 161 and 179 corresponding to hexosyl or inositol residue. We also detected characteristic fragment ions of uronic acids at m/z 175 [*M*-H_2_O-H]^-^ and 113 [*M*-COO]^-^. Consistent with the *m/z* 597 fragmentation mechanism, the fragment ion at *m/z* 293 is formed by the loss of a hexose or an inositol and a further decarboxylation. We conclude that *m/z* 517 likely corresponds to the non-phosphorylated form of the *m/z* 597 candidate.

**Figure 2.**
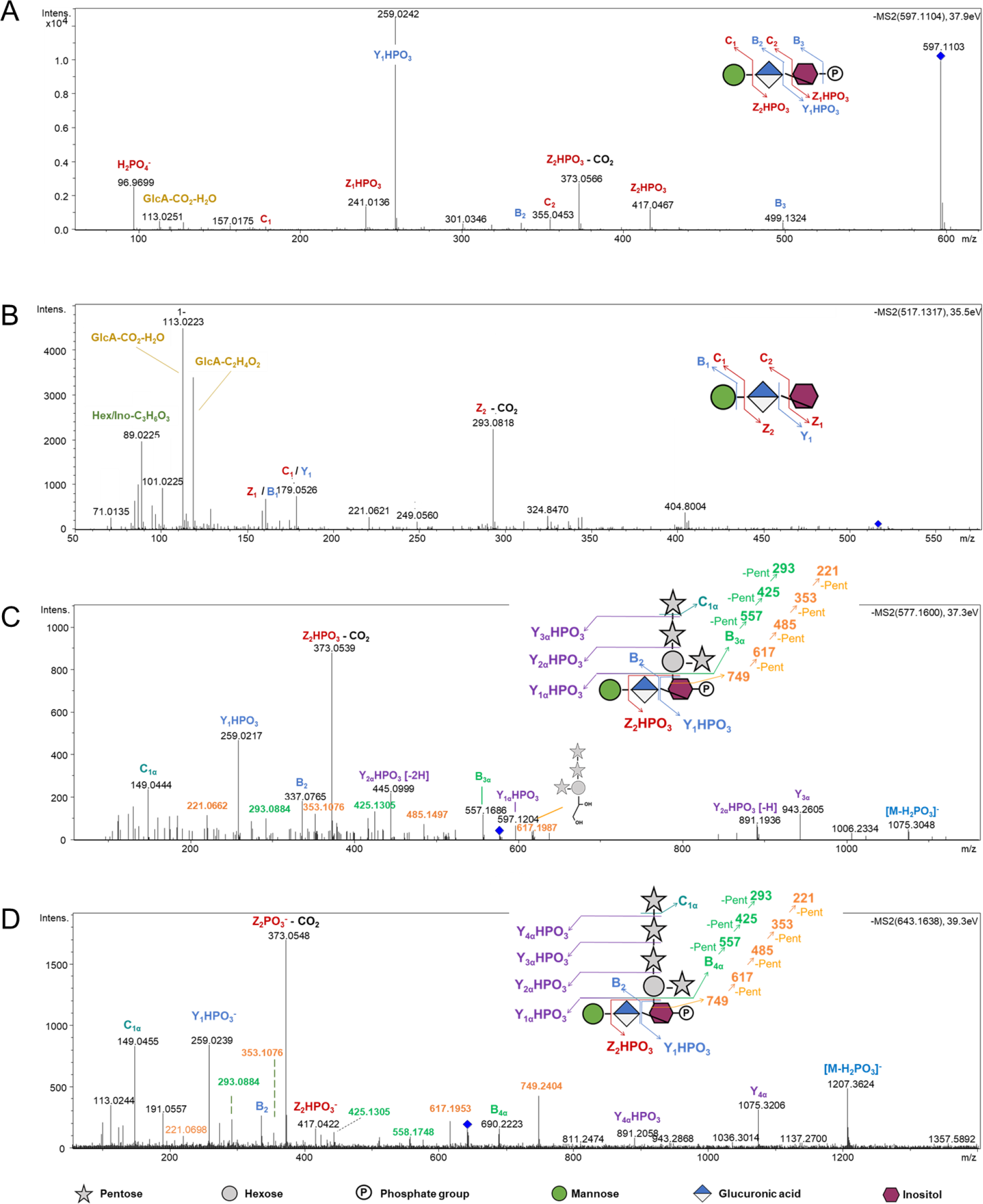
Inositol(Phosphate) Glycans (I(P)Gs) accumulate during Arabidopsis thaliana-Botrytis cinerea infection. (**A**) MS^2^ fragmentation pattern of *m/z* 597 in negative mode. (**B**) MS^2^ fragmentation pattern of m/z 517 in negative mode. (**C**) MS^2^ fragmentation pattern of m/z 577 in negative mode. (**D**) MS^2^ fragmentation pattern of m/z 643 in negative mode. GlcA: Glucuronic acid, Intens.: signal intensity. Pent: Pentose.

The *m/z* 259 (Ino-P) and 373 (UA-Ino-P) ions observed in *m/z* 597 MS^2^ spectrum were found in the fragmentation pattern of six doubly charged candidates at m/z 577 (C_39_H_63_O_37_P^2-^), 643 (C_44_H_71_O_41_P^2-^), 716 (C_50_H_81_O_45_P^2-^), 724 (C_50_H_81_O_46_P^2-^), 782 (C_55_H_89_O_49_P^2-^), and 790 (C_55_H_89_O_50_P^2-^). According to the fragmentation pattern obtained from *m/z* 577 and 643 (**Fig. 2C,D**), these molecules contain one additional hexose and three and four additional pentoses respectively, likely linked to a Hex-UA-Ino-P core. The cross-ring fragments at m/z 353, 485, 617 and 749 (B_4α_ + C_2_H_4_O_2_) might be produced by the cleavage of an inositol or a hexose residue. We favor the first hypothesis since no fragment corresponding to three or four pentoses plus two hexoses has been detected. Last, the cross-ring fragment at *m/z* 221 (C_6_H_10_O_5_ + C_2_H_4_O_2_) provides evidence that the Pent_4_Hex chain and Pent_3_Hex are branched on the inositol moiety via the hexose residue. We conclude that the *m/z* 577 and 643 candidates were IPGs.

Following the same reasoning, according to the fragmentation pattern obtained from candidates at *m/z* 716 and 724 (**Fig. S2A and S2B**), we attributed the respective fragments at *m/z* 895 and 911 to the presence of side chains composed of one hexose linked to the Hex-UA-Ino-P core and four pentoses and one additional deoxyhexose (*m/z* 716) or hexose (*m/z* 724). We finally deduced that the candidates at *m/z* 782 and 790 are characterized by one additional pentose (**Fig. S2C and S2D**). Therefore, the candidates at *m/z* 716 and 724 on one hand, and *m/z* 782 and 790 on the other hand were also IPGs.

### Inositol phosphate glycans originate from plant sphingolipids

To explore whether the I(P)Gs mentioned above originate from the plant sphingolipids known as glycosyl inositol phosphoryl ceramides (GIPCs), we analyzed the cell wall water extract of *gmt*.*1*.*3* (GIPC Mannosyl Transferase) mutant which is impaired in GIPC glycan biosynthesis (*18*). We examined 14-day-old light-grown wild-type and *gmt1*.*3* plants that exhibited similar heights. In cell wall water extract from WT plantlets, we detected six of the IPGs and IG that accumulated upon infection (**Fig. 3A,B**). In contrast, none of the oligosaccharides mentioned earlier were found in the *gmt1*.*3* mutant. We observed ions at *m/z* 435 (C_12_H_20_O_12_P) and 355 (C_12_H_20_O_12_) which were absent in the WT samples (**Fig. 3A,B**) and could correspond to GlcA-Ino-P and GlcA-Ino, respectively. The fragmentation of *m/z* 435 generated ions at *m/z* 259, likely resulting from the loss of an uronic acid, as well as ions at *m/z* 97 and 79, indicative of phosphorylated groups (**Fig. 3C**). The fragmentation of m/z 355 suggested the presence of a C_6_H_10_O_6_ moiety and an uronic acid not present in the WT samples, which we attributed to GlcA-Ino (**Fig. 3D**). It is worth to note that we did not detect any ion corresponding to the GlcA-Ino core substituted by side chains, raising question about the GIPC glycan biosynthetic machinery. In conclusions, we deduced that the seven I(P)Gs that accumulated during *A. thaliana/B. cinerea* interaction derives from Series A and Series E to H GIPCs and are likely released by GIPC phospholipase(s) C that cleaves between the phosphate and the lipid moiety (**Fig. 3D**).

**Figure 3.**
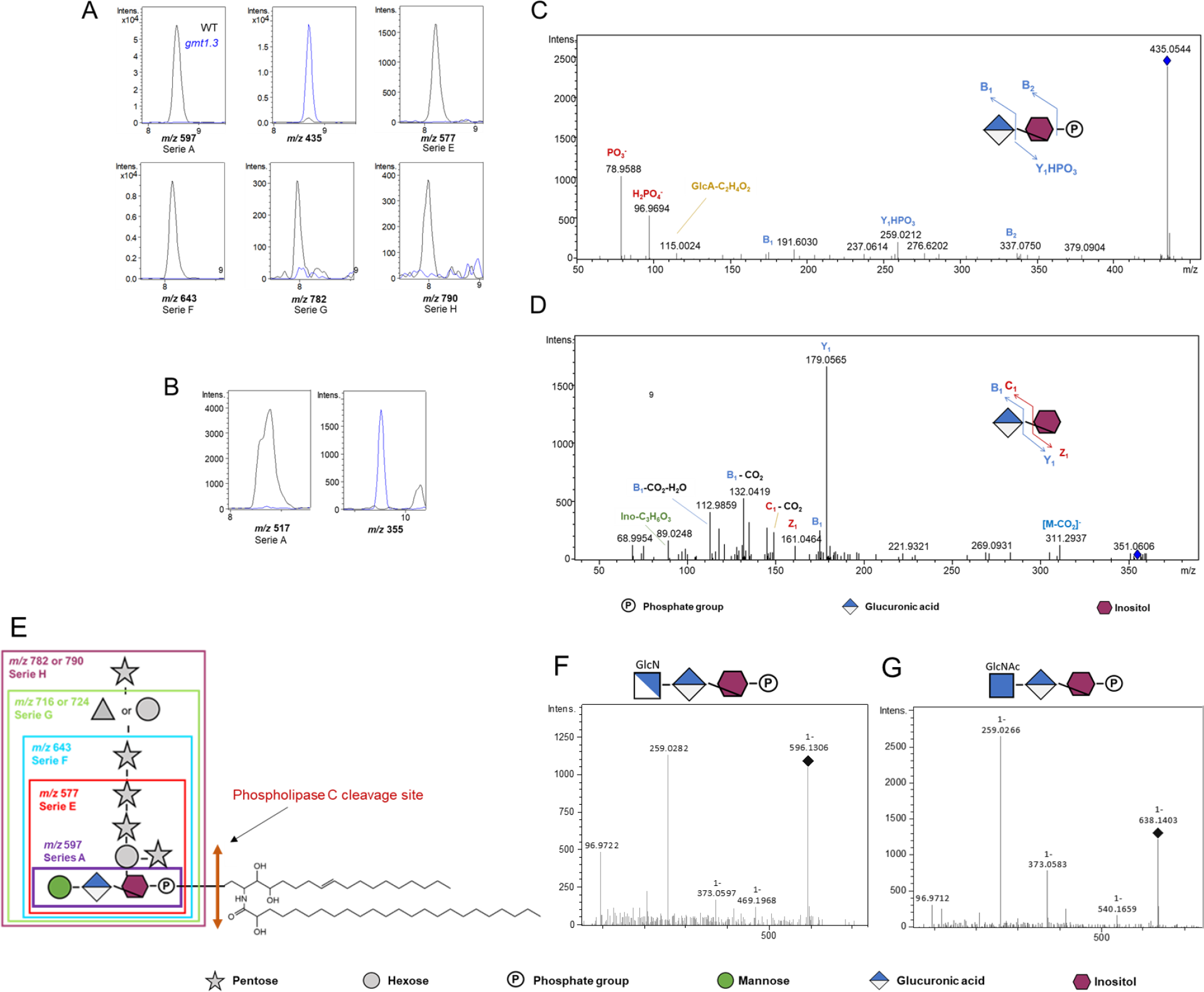
Inositol(Phosphate) Glycans (I(P)Gs) derives from plant glycosylinositol phosphorylceramides (GIPCs) Extracted ion chromatograms of IPG (**A**) and IG (**B**) in cell wall water extract of WT and *gmt1*.*3 A. thaliana* 14*-*day-old plantlets. MS^2^ fragmentation pattern of m/z (**C**) 435 and (**D**) 355, detected in *gmt1*.*3* mutant. (**E**) Structure of the different GIPC Series. Fragmentation pattern of m/z (**F**) 596 and (**G**) 608 detected in tomato leaves infected by *B. cinerea*. Intens.: signal intensity. GlcN(Ac): (N-acetyl)glucosamine.

Additionally, as the polar heads of tomato GIPC contain an acetylglucosamine residue instead of mannose (*19*), we aimed to investigate whether we could consistently detect the production of Series A IPG. We observed an ion at m/z 596 [M - H]-, for which the software predicted the formula C_18_H_31_O_19_NP^-^ and another at 638 (C_20_H_33_O_20_NP^-^). The variance in molecular formula between these two ions corresponds to an acetyl group (C_2_H_2_O). Upon fragmentation of these ions, we observed the formation of ions at m/z 259 and 373, which are characteristic of the GIPC polar (*16, 17*; **Fig. 3E,F**). We concluded that the ions at m/z 596 and 638 correspond to Series A and acetylated Series A GIPC polar heads, respectively.

### *B. cinerea* secretes phospholipases responsible for the production of Inositol Phosphate Glycans

Subsequently, in order to determine whether these IPGs were generated by the enzymatic activities of *B. cinerea* rather than originating from the plant itself, we incubated *B. cinerea* spore solutions with inert cell wall material extracted from *A. thaliana* leaves and monitored IPG production over time. Our observations revealed the gradual accumulation of IPGs from Series A to H (**Fig. 4A**), whereas none of them were detected when *B. cinerea* spores were cultivated on pectins (data not presented). This finding confirms that the production of IPGs is indeed a result of the degradation of plant GIPCs by one or more *B. cinerea* phospholipases. We therefore selected, from published transcriptome profiles (*20*), the most abundant transcript, encoding a putative sphingomyelinase in germinating *B. cinerea* conidia (*BcGIPC-PLC1*) and we cloned the CDS of this sphingomyelinase including HIS tag in order to express the enzyme in *P. pastoris*. After methanol induction, the enzyme, was secreted in the culture medium (**Fig. S3**) and showed GIPC-PLC activity on Series A GIPCs and to a lesser extent on Series F, G and H (**Fig. 4B**) and a preference for acidic pH (**Fig. 4B**). This suggests that *B. cinerea* secretes other GIPC-PLCs with varying substrate specificities.

**Figure 4.**
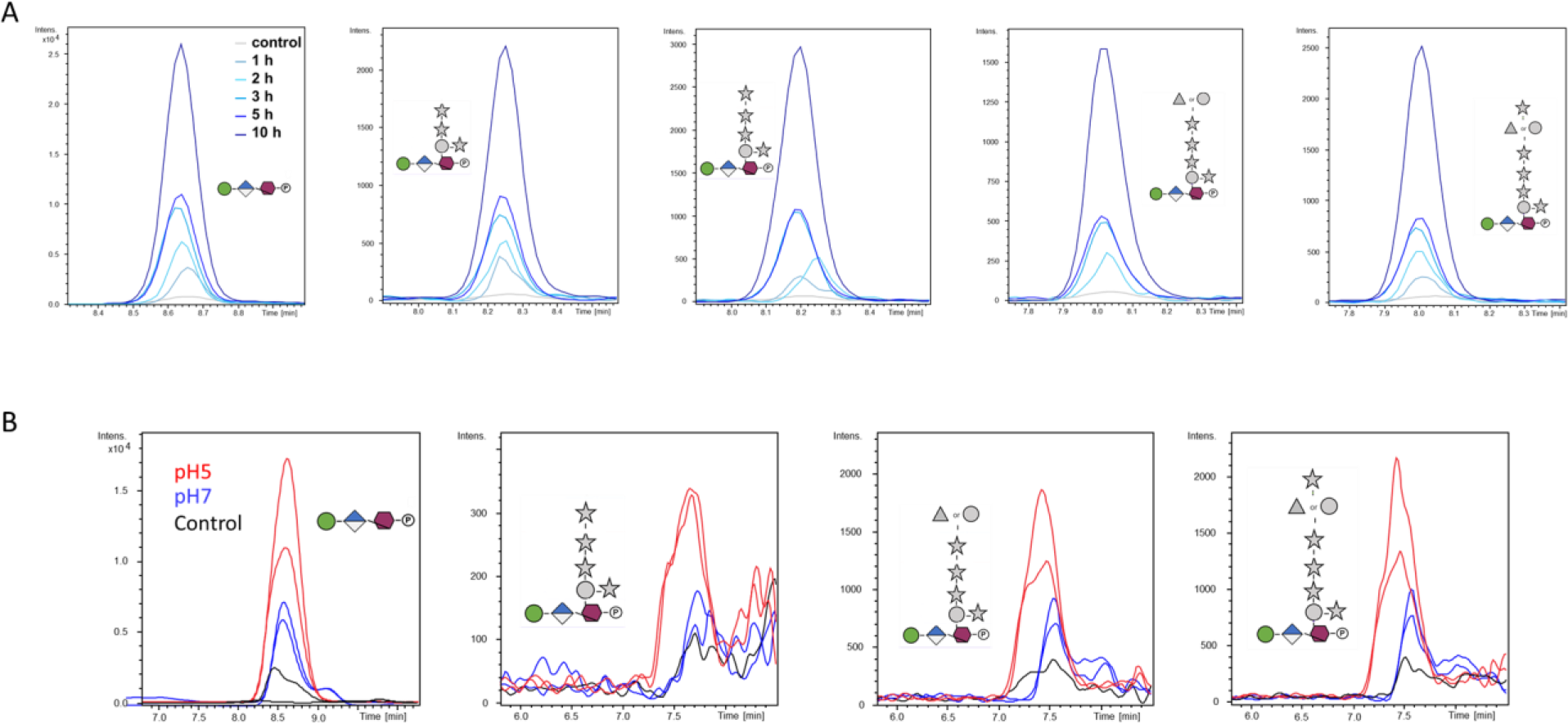
*B. cinerea* is responsible for the production of InositolPhosphate Glycans (IPGs) from plant membrane upon infection. **(A)** Extracted ion chromatograms of accumulated IPGs over time. *B. cinerea* spores were incubated with alcohol-insoluble residues prepared from *A. thaliana* leaves, and the production of each IPG was monitored using HPSEC-HRMS at 1, 2, 3, 5 and 10 hours (light blue to dark blue). (**B**) Extracted ion chromatograms of IPGs produced by the phospholipase encoded by Bcin07g04350. The enzyme was expressed in an inducible recombinant protein production in *Pichia pastoris*. Black line: culture media from non-induced cultures incubated with alcohol-insoluble residue containing plant sphingolipids at pH 5. Red and blue: culture media from methanol-induced cultures incubated with alcohol-insoluble residue containing plant sphingolipids at pH 5 and 7, respectively.

Finally, to determine whether some of these IPGs could function as damage-associated molecular pattern, we purified Series G and H IPGs from cell wall water extract of *A. thaliana* 5-week-old leaves using size-exclusion chromatography. We assessed the purity of collected and dialyzed fractions by HPSEC-HRMS. We used the promoter of JASMONATE-ZIM-DOMAIN PROTEIN 10 gene (21,22) and MYB DOMAIN PROTEIN 51 (23) gene to drive the transcription of a secretable α-glucuronidase (GUS) based reporter as tools to monitor the expression of negative regulators of jasmonic acid (JA) and indolic glucosinate biosynthesis, respectively, involved in plant defense during *B. cinerea* infection. When we infiltrated *A. thaliana* leaves with a solution containing Series G and H IPGs at a concentration of 40 µg/mL, we observed significant increase in the expression of *JAZ10* gene using quantitative and qualitative GUS activities (**Fig. 5A,C**). Interestingly, Series G and H IPGs also led to a reduced expression of a reporter gene associated with indolic glucosinolate biosynthesis (**Fig. 5B,D**) This suggests that Series G and H IPGs tend to suppress plant defenses rather than activate them. To verify this hypothesis, we infiltrated *A. thaliana* leaves with this solution of Series G and H IPGs and infected these leaves with *B. cinerea* 8 hours later. We observed, 72 hours post infection, that plant infiltrated with IPGs exhibited increased sensitivity to *B. cinerea*, confirming that Series G and H IPGs have the ability to downregulate the plant defense mechanisms (**Fig. 5E**).

**Figure 5.**
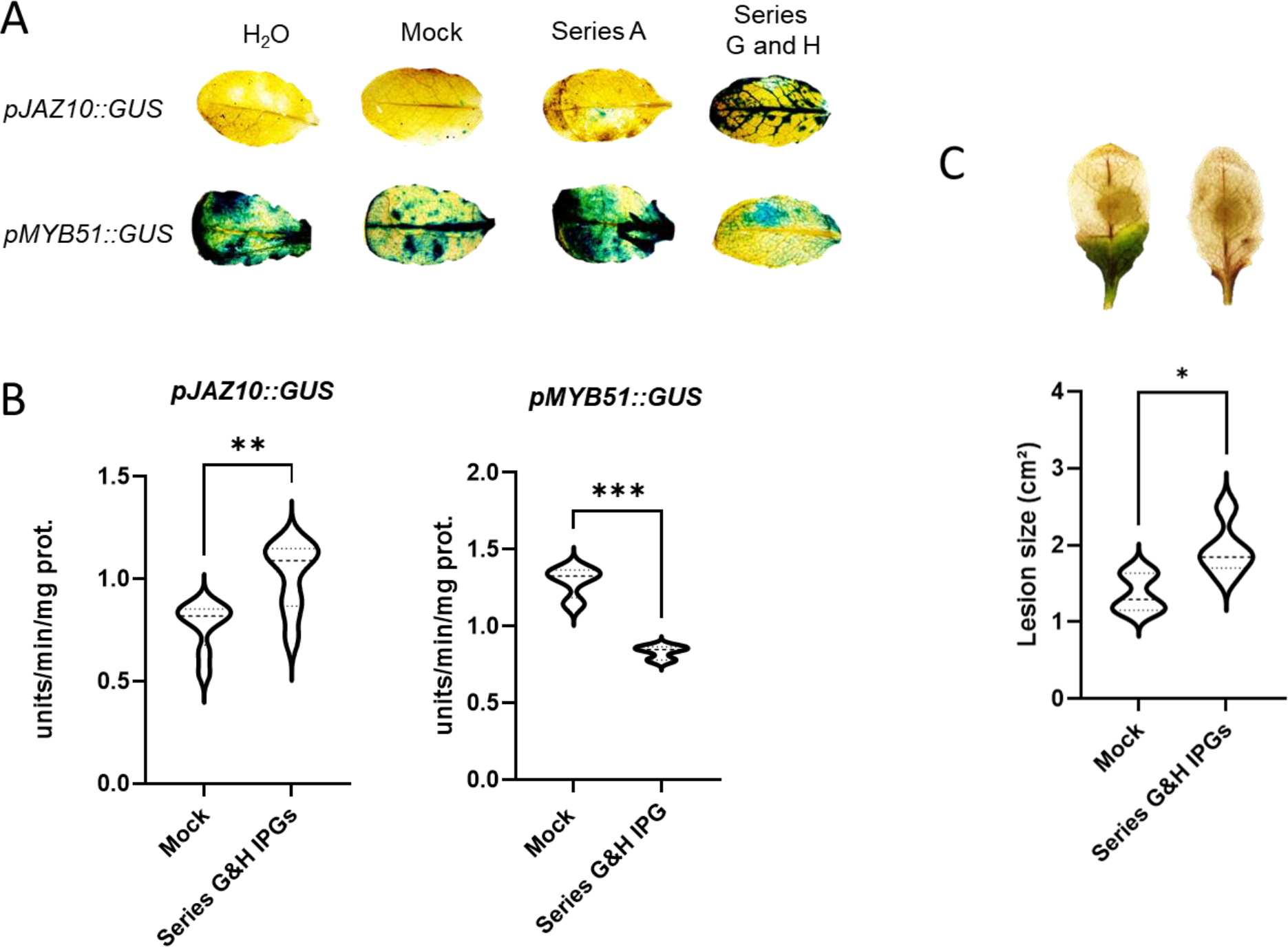
Series G and H InositolPhosphate Glycans downregulate plant defense. **(A)** GUS staining of *A. thaliana* leaves expressing the promoter JAZ10::GUS fusion or MYB51::GUS fusion after infiltration of Series A or G and H IPGs **(B)** GUS activity quantified by fluorescence in 5-week-old *A. thaliana* rosette leaves expressing the promoter *JAZ10::GUS* or *MYB51::GUS* fusion after the infiltration of Series G and H IPGs. Values are means ± SD (n = 4); **P < 0.01 and ***P < 0.001 by Student test. (C) Infection assays of *B. cinerea* on 5-week-old *A. thaliana* rosette. Leaves were filtrated with a mixture of Series G and H IPGs and were infected 8 hours later by *B. cinerea*. Lesion size of the infected leaves were measured 72 h later. Values are means ± SD (n = 6); *P < 0.05 by Student test.

## Discussion

We unveiled the presence of previously undiscovered oligosaccharides that accumulate during the interaction between plants and *B. cinerea*. Despite the anticipation of characterizing pectin-derived oligosaccharides, our study revealed the prominent presence of xyloglucan fragments and IPGs originating GIPCs upon infection.

In plants, one IG has already been identified in plant culture cells (24, 25). However, to our knowledge, our study represents the first report of IPGs in plants, while these compounds have been well-documented in mammals for over 30 years (*26*). We demonstrated the presence of IPGs from Series A to H in cell wall extracts of healthy plants, which are preferentially accumulated upon infection. The fact that *B. cinerea* was able to produce these IPGs from inert plant material prompted our investigations into the existence of genes encoding GIPC-PLC into *B. cinerea* genome. We successfully identified a GIPC-phospholipase in *B. cinerea*, annotated as a sphingomyelinase, predominantly active on Series A GIPCs, indicating its potential role in the generation of these IPGs during the interaction between plants and *B. cinerea*.

It is worth to note that the non-specific phospholipase C4 (NPC4) in *A. thaliana* has also been shown to be involved in GIPC hydrolysis in response to phosphate deficiency (*27*). This finding is consistent with the fact that we also detect IPGs in non-infected leaves. The accumulation of series A inositol glucan (*m/z* 517) also suggests that a GIPC phospholipase D, which cleaves after the phosphate moiety, could be activated. A phospholipase D activity has been detected in cabbage and in *A. thaliana* (*28*). This GIPC-PLD prefers GIPC containing two sugars (*29*) which aligns with the fact that Series A is the most highly produced in our conditions. However, we cannot rule out another scenario that would imply a phospholipase C activity and a subsequent phosphorylase during infection.

Furthermore, the absence of series E to H IPGs in the *gmt1*.*3* mutant, which lacks mannose residues, suggests that mannose addition is indispensable before the transfer of other sugars. Notably, this finding raises questions about the role of complex GIPC lack in the *gmt1*.*3* mutant’s severe phenotype and constitutive hypersensitive response (*18*). The potential role of IPGs as suggesting that IPGs could act more broadly as stress-regulator molecules. signaling molecules warrants further investigation in this mutant and across the plant kingdom

Our research also uncovers the vital finding that certain IPGs, specifically those from Series G and H, have the ability to downregulate the plant’s defense mechanisms. Altogether these data suggest that IPGs could act in non-infected plant as defense-regulator molecules and that *B. cinerea* has evolved to exploit this signaling pathway to its advantage. This discovery underscores the fungus’s production of molecules capable of circumventing plant defense and sheds light on the dynamic strategies employed by *B. cinerea* to undermine the plant’s protective responses and establish a successful infection and further highlight the dynamic interactions between the plant and the pathogen

Finally, it is worth noting that bacterial sphingomyelinases, which enzymatically break down sphingomyelin in mammals, release products identical to those generated by host eukaryotic enzymes that aids in evasion of the immune response (30). Therefore, our study provides unique insights into shared strategies that plant and mammal pathogens use to infect and overcome host defenses. This can help identify common vulnerabilities that could be targeted for disease control in both contexts.

## Acknowledgments

J. Mortimer, M-C Soulié and D. Gasperini are thanked for providing *gmt1* seeds, B05.10 strain and pJAZ10::GUS and pMYB51::GUS transformants, respectively. This work has benefited from the support of IJPB’s Plant Observatory technological platforms.

## Author contributions

Conceptualization: AV, SV

Methodology: AV, SV, TD

Investigation: AV, LL, JR, JS, EA, AF

Funding acquisition: AV, SV

Project administration: SV

Supervision: AV, SV

Writing – original draft: AV

Writing – review & editing: AV, SV, JS

## Data and materials availability

All data are available in the main text or the supplementary materials.

