## supplemental material for "Inositolphosphate glycans accumulate and suppress plant defense during Arabidopsis/Botrytis interaction"

### Materials and Methods

Plant Material and growth

Infection assays were performed on *A. thaliana* WT Wassilewskija (Ws-0) plants grown in soil in a growth chamber at 22 °C, 70% humidity, under irradiance of 100 mol.m$^{-2}$.s$^{-1}$ with a photoperiod of 8 h light/16 h dark. Seeds from *gmt1.3* mutants were a kind gift of J. Mortimer. pJAZ10::GUS and pMYB51::GUS transformants were a kind gift of D. Gasperini. *A. thaliana gmt1.3* and Colombia seeds were grown on 1/2 × MS media plates positioned vertically for 14 days under constant light at 23 °C. The tomato *Solanum lycopersicum* was cultivated in greenhouse conditions in soil.

The *B. cinerea* B05.10 collected from Vitis in Germany (*31*) was used as the wild-type (WT) reference strain. It was grown on potato dextrose agar at 23°C under continuous light.

Alcohol insoluble residue and cell wall water extract

Leaves and plantlets were grinded in five volume of ethanol 96% and the supernatant was removed after centrifugation at 10,000*g* for 5 min. The pellet was washed with 70% ethanol with subsequent centrifugation until being uncolored and dried in a speed vacuum concentrator at room temperature. Two volumes of distilled water were added to the alcohol insoluble residue obtained and leave one hour at room temperature. The supernatant was collected after centrifugation at 10,000 *g* for 5 min and dried in a speed vacuum concentrator at room temperature.

Oligosaccharides accumulated upon *B. cinerea* infection of *A. thaliana* and *S. lycopersicum* leaves

Oligosaccharides were produced and analyzed according to Voxeur et al., 2019 (*13*).

Briefly, after 10 days, the spores of *B.cinerea* were washed from the surface of the plate using Gamborg's B5 basal medium, 2% (w/v) fructose and 10 mM phosphate buffer. Fungal hyphae were removed by filtering. The concentration of spores was determined using a Malassez cell and adjusted to a final concentration of 3.10$^5$ conidia/mL. Isolated tomato leaves of 4-week-old plants or *A. thaliana* leaves of 5-week-old plants were immersed in a *B. cinerea* suspension (6 leaves for 10 ml of suspension at 3 x 10$^5$ spores/ml) and incubated on a rotary shaker at 100 rpm at 23 °C during 12, 15 and 18 h. The liquid medium was next collected and an equal volume of 96% ethanol was added. After centrifugation at 5000 *g* during 10 min, the supernatant was collected and dried in a speed vacuum concentrator at room temperature. The obtained pellet was then diluted. The equivalent of the digestate of 3 leaves of 5-week-old *A. thaliana* plants and similar leave surface of tomato was dried and diluted in 200 µl. 10 µl were injected for MS analysis.

Oligosaccharide analysis

Samples were diluted at 1 mg/ml in ammonium formate 50 mM, formic acid 0.1%. Chromatographic separation on high-performance size-exclusion chromatography was performed on an ACQUITY UPLC Protein BEH SEC Column (125Å, 1.7 µm, 4.6 mm x 300 mm, Waters Corporation, Milford, MA, USA). Elution was performed in 50 mM ammonium formate, formic acid 0.1% at a flow rate of 400 µl/min and a column oven temperature of 40 °C. The injection volume was set to 10 µl. MS-detection was performed on a Bruker impact II QTOF in negative mode with the end plate offset set voltage to 500 V, capillary voltage to 4000 V, Nebulizer 40 psi, dry gas 8 l/min and dry temperature 180 °C. Major peaks were annotated following accurate mass annotation, isotopic pattern and MS/MS analysis. The MS fragmentation pattern is indicated according to the nomenclature of Domon and Costello (*32*). For the targeted analysis, the theoretical exact masses were used with 4 significant figures with a scan width of 5 ppm. The resulting extracted ion chromatograms were integrated.

Data processing

The .d data files (Bruker Daltonics, Bremen, Germany) were converted to .mzXML format using the MSConvert software (ProteoWizard package 3.0 (*33*)) mzXML data processing, mass detection, chromatogram building, deconvolution, sample alignment, and data export were performed using MZmine 2.52 software (http://m mzmine.github.io/). Mass list were built using a retention time window of 6.0-8.0 min and next, we used the ADAP chromatogram builder (*34*) with group size of scan 5, peak detection threshold of 800, a minimum highest intensity of 1500 and *m/z* tolerance of 0.01 m/z. We deconvoluted the data with the ADAP wavelets algorithm using the following setting: S/N threshold 8, peak duration range = 0.01-0.2 min RT wavelet range 0.02-0.1 min. Finally, we selected candidates with a surface area superior to 18000 at 18 hpi and results were deisotoped manually.

Isolation of IPGs

100 mg of freeze-dried 30-day-old leaves was suspended in 15 ml of aqueous 70% (v/v) ethanol overnight at room temperature. The supernatant was discarded and after drying, the residue was grinded using metal balls. The alcohol insoluble residue obtained treated with 1 mL of distilled water during two hours and the supernatant was collected to purify IPGs. The IPGs were separated other solubilized molecules by size-exclusion chromatography on a HiLoad™ 16/600 Superdex™ 30 pg column (Cytiva) in 200 mM sodium acetate at pH5 at a flow rate of 1mL/min and detected with a refractive index detector (Refractomax, Thermofisher). Fractions were dialyzed (100 Da cutoff dialysis tubing) against deionized water and freeze-dried.

*BcGIPC-PLC* cloning and enzymatic activities.

The CDS of the putative sphingomyelinase *BCIN07g04350* from *B. cinerea* (21-630 aa) was cloned in frame with the □ factor sequence allowing the protein secretion and the C terminal peptide containing the *c-myc* and 6XHis- epitopes into the vector pPICZ□A (Proteogenix, https://www.proteogenix.science/fr). Codon optimization was used to improve the expression of the protein in *Pichia pastoris X33*. The yeast strain was transformed with 10 □g of the linearized plasmid digested by *SacI* enzyme according to the easyselect yeast expression kit (Invitrogen). Transformants were isolated and analyzed for the presence of the insert using *AOX1* primers according to the easyselect yeast expression kit (Invitrogen). The successfully obtained transformants were grown in baffled flasks in 1.5 mL of buffered glycerol-complex medium, overnight at 30°C using 100 mg/ml Zeocin. Cells were then collected by centrifugation and resuspended to an OD600 of 1.0 in 100 mL of buffered methanol complex medium. A final concentration of 0.5 % (v/v) methanol was added every 24 h to maintain induction. After 72 h of induction, the culture was centrifuged at 1 500 g for 10 min and the culture supernatant was loaded on Amicon ultra-4ml -30 kDa cutoff (Merck millipore). To identify the recombinant protein present in the supernatant by Western blot, SDS-PAGE was transferred from resolving gel to PVDF blotting membrane using the appropriate cathode and anode buffers and a Trans-Blot TURBO Transfer System (Bio-Rad, Cat. No. 170-4155) at 120V for 60 min. TBS-T (0.5 % Tween 20 in TBS) was used as washing buffer and 5% non-fat dried milk in TBS-T was used as blocking reagent. Transferred proteins were incubated for 1 h at room temperature under shaking with 1:3000 dilution of anti-his antibody coupled with peroxidase (Sigma, Cat. No. A7058). After washes, the ECL kit (Cytiva) was used to detect the protein of interest according to the supplier's instructions. 200 □l of the concentrated supernatants of induced and non-induced cell culture were next mixed with 10 mg of alcohol insoluble residues obtained from *A. thaliana* 5-week-old overnight at 37 °C at pH5 or 7.

Table S1: Primers AOX1

| | |
|---|---|
| 5' *AOX1* primer | 5'-GAC TGG TTC CAA TTG ACA AGC-3' |
| 3' *AOX1* primer | 5'-GCA AAT GGC ATT CTG ACA TCC-3' |

**Qualitative and quantitative GUS activities.**

A solution of 40 mg/ml of series H and G IPG fractions were next infiltrated in leaves of 5-week-old seedlings expressing the following reporter constructs *pJAZ10::GUS.* GUS or *pMYC51::GUS*. To perform qualitative GUS assays, leaves were submerged in GUS buffer as described (*35*) and infiltrated three times (1 min) under vacuum, and incubated at 37°C for 5h. Seedlings were washed three times with 70% ethanol.
Quantitative activity analyses were performed on the aerial part of 5-week-old seedlings as described (*36*) with some modifications: the GUS buffer does not contain any β-mercaptoethanol and the measures were performed with a fluoroskan ascent (Thermo Scientific, Waltham, MA, USA). Three pools of two leaves of two different replicates were analyzed.
GUS staining

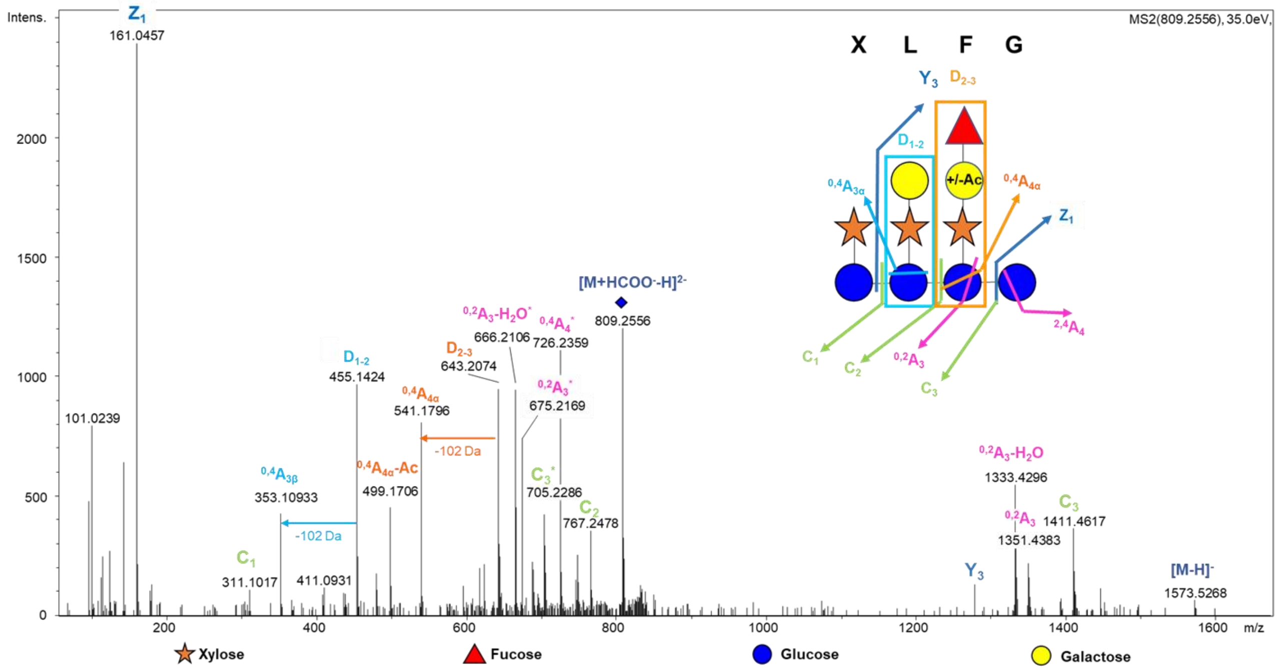


**Fig. S1. An acetylated and fucosylated xyloglucan oligosaccharide is accumulated upon *Arabidopsis thaliana-Botrytis cinerea* infection**. $MS^2$ fragmentation pattern of *m/z* 809 and a proposed fragmentation scheme.

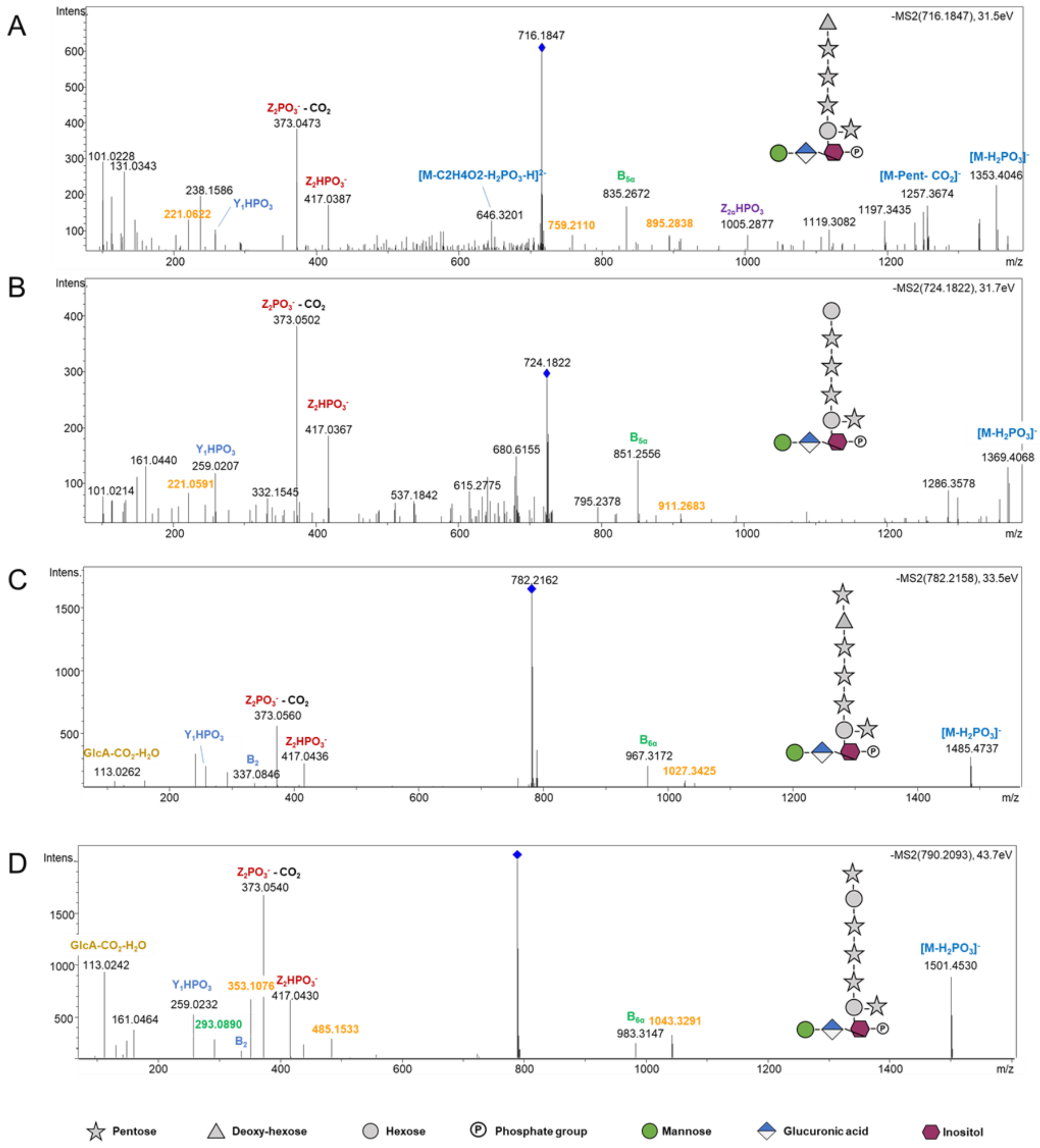

**Fig. S2. Inositol(Phosphate) Glycans (I(P)Gs) accumulate during *Arabidopsis thaliana-Botrytis cinerea* infection** $MS^2$ fragmentation pattern of *m/z* (**A**) 716, (**B**) 724, (**C**) 782 and (**D**) 790 in negative mode. GlcA: Glucuronic acid, Intens.: signal intensity.

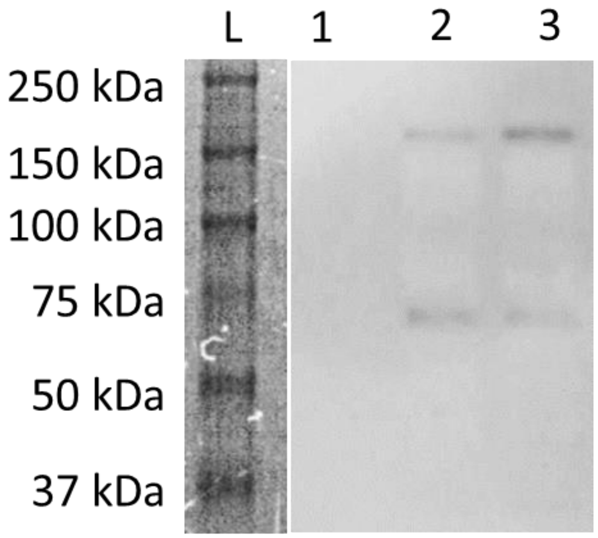


**Fig. S3. Secretion of the BcGIPC-PLC (68.5 kDa expected weight) encoded by *BCIN_07g04350* in the X33 *Pichia pastoris* strain** Anti-His antibodies and chromogenic detection were employed. L: molecular weight markers with an acquisition 17 min. 1: Proteins extracts from the supernatant of *BcGIPC-PLC* transformant culture non-induced. 2. Protein extracts from the supernatant of *BcGIPC-PLC* transformant culture n°1 induced with methanol during 72 h. 3 Protein extracts from the supernatant of *BcGIPC-PLC* transformant culture n°2 induced with methanol during 72 h. Two min of acquisition were used to reveal the His-tagged expressed proteins.

**Data S1. (separate file)**

Retention time, mass-to-charge ratio and peak area of candidates.

**References.**

31. T. Quidde, A. E. Osbourn, P. Tudzynski. Detoxification of α-tomatine byBotrytis cinerea. *Physiological and Molecular Plant Pathology*, *52*(3), 151-165. (1998). doi: 10.1006/pmpp.1998.0142

32. B. Domon, C. E. Costello. Systematic Nomenclature for Carbohydrate Fragmentations in FAB-MS/MS Spectra of Glycoconjugates. *Glycoconjugate Journal*, 5 (4), 397–409. 28 (1988). doi: 10.1007/BF01049915

33. M. C. Chambers, B. Maclean, R. Burke, D. Amodei, D. L. Ruderman, S. Neumann, L. Gatto, B. Fischer, B. Pratt, J. Egertson, K. Hoff, D. Kessner, N. Tasman, N. Shulman, B. Frewen, T. A. Baker, M. -Y. Brusniak, C. Paulse, D. Creasy, L. Flashner, K. Kani, C. Moulding, S. L. Seymour, L. M. Nuwaysir, B. Lefebvre, F. Kuhlmann, J. Roark, P. Rainer, S. Detlev, T. Hemenway, A. Huhmer, J. Langridge, B. Connolly, T. Chadick, K. Holly, J. Eckels, E. W. Deutsch, R. L. Moritz, J. E. Katz, D. B. Agus, M. MacCoss, D. L. Tabb, P. A. Mallick. A Cross-Platform Toolkit for Mass Spectrometry and Proteomics. *Nature Biotechnology*, 30 (10), 918–920. 29 (2012). doi: 10.1038/nbt.2377

34. O. D. Myers, S. J. Sumner, S. Li, S. Barnes, X. Du. One Step Forward for Reducing False Positive and False Negative Compound Identifications from Mass Spectrometry Metabolomics Data: New Algorithms for Constructing Extracted Ion Chromatograms and Detecting Chromatographic Peaks. *Analytical chemistry*, 89(17), 8696–8703 (2017) doi:10.1021/acs.analchem.7b00947

35. R. A. Jefferson, T. A., Kavanagh, M. W. Bevan. GUS fusions: beta-glucuronidase as a sensitive and versatile gene fusion marker in higher plants. *The EMBO journal*, *6*(13), 3901–3907 (1987). doi:10.1002/j.1460-2075.1987.tb02730.x

36. T. Elmayan, H. Vaucheret. Expression of single copies of a strongly expressed 35S transgene can be silenced post-transcriptionally. *The Plant Journal*, *9*(6), 787-797 (1996). doi: 10.1046/j.1365-313X.1996.9060787.x
